# Cellular structure analysis based on magnetic induction finite element method simulations and measurements

**DOI:** 10.1101/275271

**Authors:** J. Tang, M. Lu, W. Yin

## Abstract

Biological samples exhibit frequency dependent spectra caused by the dispersion mechanism which describes a phenomenon of dielectric relaxation due to the interaction between electromagnetic field and biological samples at cellular levels. Changes in cellular structure such as cellular shape, membrane thickness and integrity could affect bio-impedance spectra.

In this paper, the influence of cellular shape, membrane thickness and integrity on dielectric properties of biological cells suspension is simulated using a custom developed FEM solver. In its 2D version, the AC conduction case is simulated. In its 3D version, magnetic induction case is simulated. And a new method for calculating the equivalent complex conductivity of biological cell suspension along the eddy current direction is introduced.

Membrane integrity on beta dispersion was experimentally investigated using AC conduction (contact electrode) method on potato samples.

This suggests that bio-impedance measurements could provide indication of cellular structural changes of biological samples. This could be useful for biomedical, pharmaceutical and food inspection applications.

## 1. Introduction

It has been known for decades that the complex impedance of biological samples, while measured over a range of frequencies using time varying electromagnetic field, contains information about the physical, chemical or cellular properties of the samples. The permittivity and conductivity of the biological samples are frequency-dependent and characterised by three main dielectric dispersions named *α,β* and *γ* dispersion [1]. Each dispersion refers to a specific characteristic of biological samples. Schwan [1] introduced the dispersions:

*α*-dispersion: taking place at low frequency —from dc to hundreds hertz. This dispersion is caused by the relaxation of ions surrounding at the charged cellular membrane.

*β*-dispersion: taking place at radio frequency — mainly from hundreds hertz to megahertz. This dispersion is caused by Maxwell-Wagner effect, an interfacial polarisation of cell membrane blocking the ion-flow of intra and extra cellular dielectrics.

*γ*-dispersion: taking place at microwave frequency — gigahertz region. This dispersion is caused by relaxation of free water molecules in intra and extra cellular fluid.

**Figure 1:**
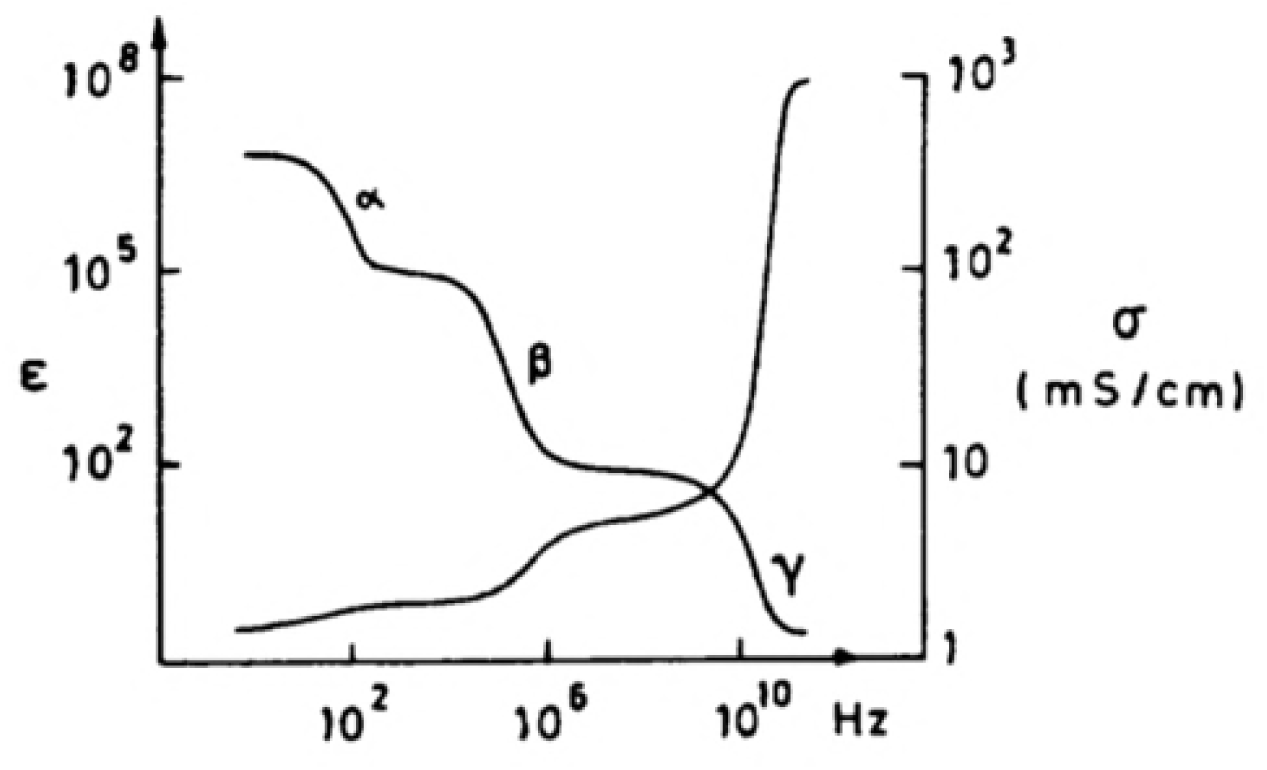
dispersions of biological samples

Theoretically, the biological samples display a specific *β*-dispersion based on the geometry structure and dielectric properties of cell membranes. The characteristics of *β*-dispersion reflect geometry and electrical information of the biological samples. Geometry structure is considered to be one of the most important peculiarities of biological samples. However, the micro-level geometry information of biological samples is hard to obtain from macro-level observations or measurements due to the small size of biological cells. For example, candida albican [12] and normal yeast are structurally different in microscopic as shown in Figure 2 while it is hard to identify them by macroscopic methods. Given that, measuring the impedance of biological samples is an effective and reliable way to distinguish the biological samples with different geometry structures and electrical properties [6][7]. Asami [2] and Marco [25] have built an analytical solution model of non-spherical biological cells to calculate the electrical properties of the cell suspensions. The axial ratio is considered to be the main factor that influences the dielectric behaviour of the non-spherical biological cell suspensions; however, they did not consider fully the structural differences such as shape, membrane thickness and integrity. In addition, most FEM simulation works about biological samples analyse the dielectric properties of bio-cell suspension by applying alternating electric field in a contact manner. For example, Sekine [3] and Asami [4] have introduced FEM simulations about spherical biological cells suspension under alternating electric field.

**Figure 2:**
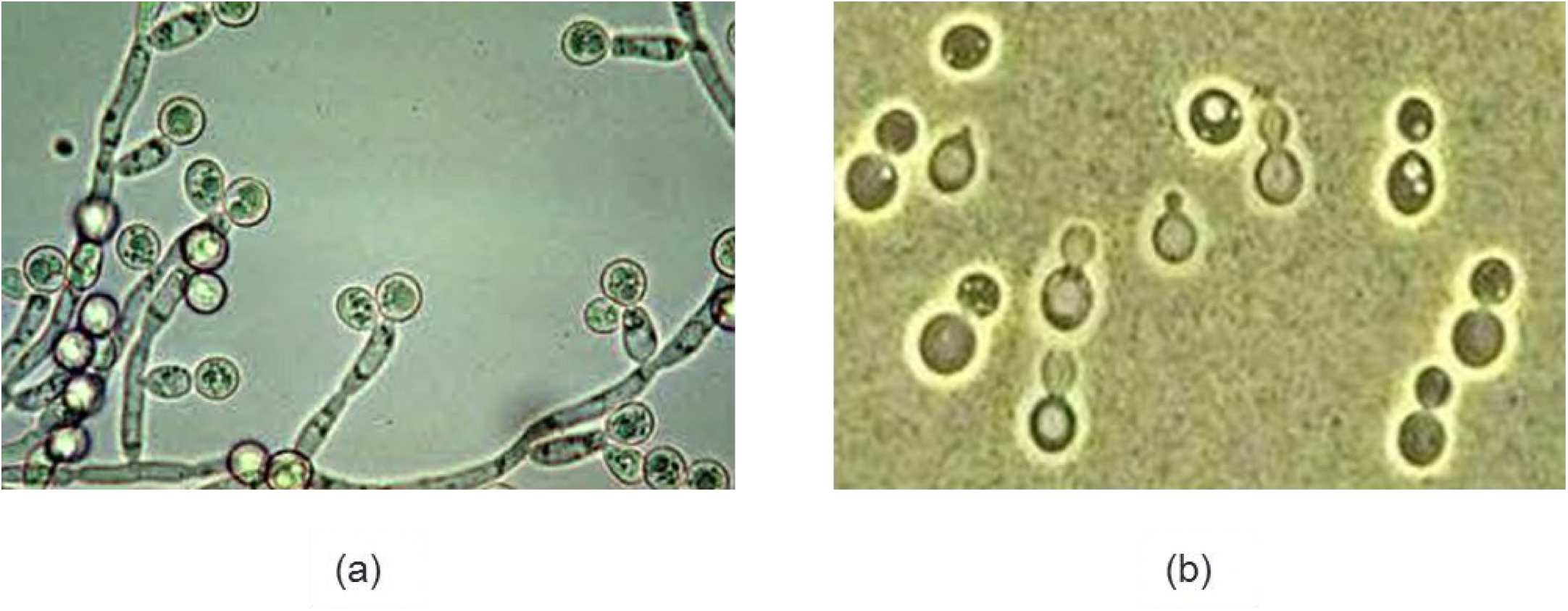
(a) Candida albican under microscope. (b) Normal yeast under microscope.

In this paper, the dielectric behaviour of bio-cells suspension is simulated by analysing eddy currents induced by alternating magnetic field in a non-contact manner. The difficulty of the FEM simulation about biological samples is that there are too many meshing elements due to the tiny thickness of biological membrane which leads to a large amount of calculation time. Given that, a Matlab FEM simulation solver has been built and the calculation time is largely reduced comparing with commercial software like COMSOL. Furthermore, a new method to calculate the equivalent conductivity and relative permittivity of the suspension in FEM eddy current simulation is introduced. These simulations can be performed in multi frequencies and therefore they are essentially virtual biological impedance spectroscopy (BIS).

The biological impedance spectroscopy (BIS) [17] [24] has been widely investigated for medical applications such as detection of tumour [8] [19], detection of cerebral stroke [9] and electroporation treatment [10]. BIS has also been used for food industry applications. For example, it has been proposed for quality inspection of meat [11], vegetables and fruits. And it has also been used to monitoring the growth of yeasts [13] and process of brewing [14].

BIS has mainly been performed by injecting current by electrodes placed on the biological samples but less so with inducing eddy currents in the biological samples by coils. The first method is to calculate the impedance of the biological samples by measuring the surface potential difference between electrodes and the current through them. This measurement method is fast and effective. However, the polarization of electrodes, which gathers charges on the surface of electrodes, produces an electrical field against the applied field and this leads to significant errors at low frequency. In this paper, we performed four-electrode measurement to minimize the errors caused by electrode polarization [18] and the measurement is simulated by two-dimension finite element method.

The second measurement method is a non-contact magnetic induction method. Eddy currents are induced in the biological samples by coils and the impedance spectroscopy is measured by detecting the resulting magnetic field / induced voltage. In this paper, 3D FEM simulations about the eddy currents in biological samples induced by coils are presented.

## 2. Background theory

### 2.1 Finite element simulation

An induction model is built for the three-dimension FEM simulation. As shown in Figure 3, the cylinder stands for the cell suspension and the four spheres are the cells. Cell models are placed randomly in the simulation in order to simulate real suspension. Imaginary transmitter and receiver coils are put on the top of the suspension to provide alternating magnetic field and measure induced secondary magnetic field and eventually obtain the equivalent conductivity and permittivity.

**Figure 3:**
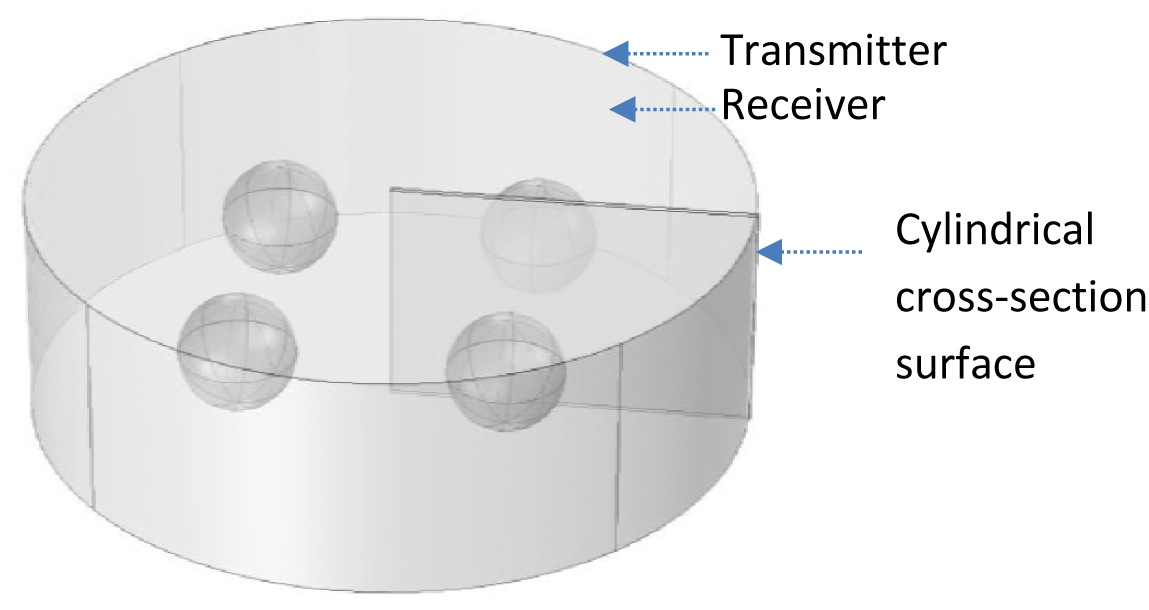
Three-dimension simulation model (4 symmetrically arranged cells)

In Figure 3, alternating current is applied to the transmitter coil and an alternating magnetic field will be induced [6][7].

The differential form of Maxwell’s equation:

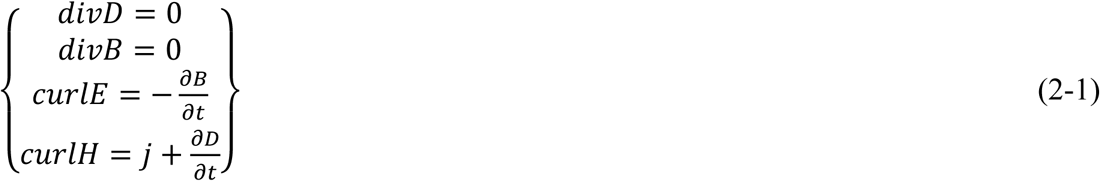

In this FEM simulation, the system is regarded to be quasi-static which assumes the influence of displacement current is very small that it could be ignored comparing with the eddy current induced by coil excitation. Time varying magnetic fields exist in nonconductive regions *Ω*_*n*_ and conductive regions *Ω*_*c*_. So the Maxwell’s equation could be written as:

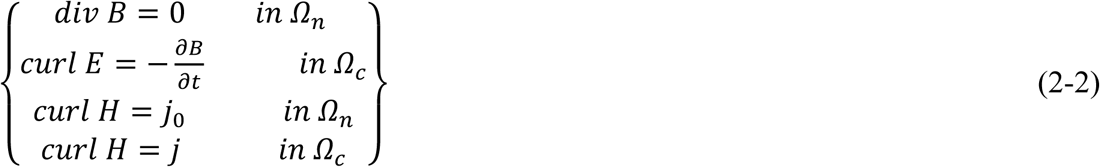

Where *j*_0_ is the given current density in *Ω*_*n*_ excited by coil. *j* is the current density in *Ω*_*c*_. H is the magnetic field intensity. B is the magnetic flux density. E is the electric field intensity.

And the field quantities are following the equations:

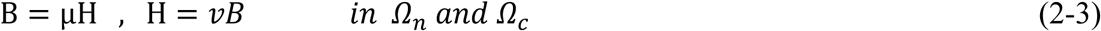

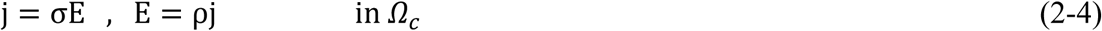

Where *μ* denotes the permeability, v is the reluctivity, σ is the conductivity and ρ is the resistivity.

According to Oszkar Biro [16], the basic formulas for the three-dimension edge elements simulation of eddy current problems, Galerkin’s equation is shown as following:

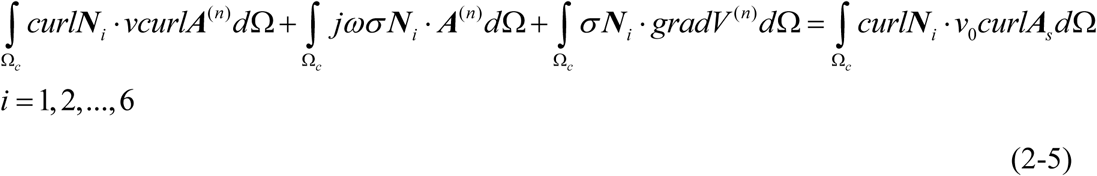

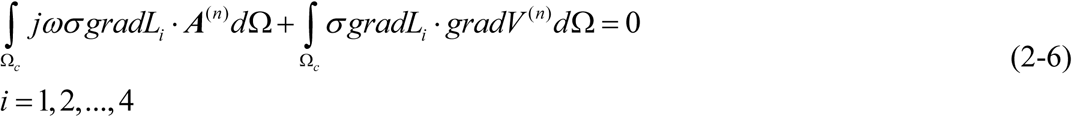

Where *N*_*i*_ is the vector interpolation of *i*^*th*^ edge corresponding to its n^th^ edge element; *L*_*i*_ represents the elemental interpolation of *i*^*th*^ node corresponding to its n^th^ element; *A*_*s*_ is the original edge vector potential of the n^th^ element;*A*^(*n*)^ is the produced edge vector potential of the n^th^ element; *V*^(*n*)^ is the electrical potential on the received (pick- up) coil contributed by the n^th^ element;*v* is the reluctivity (the reciprocal of the permeability) of the target; *v*_0_ is the reluctivity (the reciprocal of the permeability) of the air;*σ* is the conductivity of the target

The first component of equation (2-5) 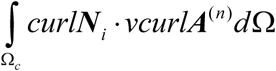 stands for the fundamental formation of the vector potential. The second term of (2-5) 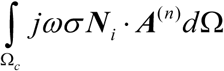 describes the diffusion effect. The third term of the equation 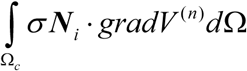 relates to Maxwell-Wagner effect. The Maxwell-Wagner effect occurs on the surface of inhomogeneous materials, so the third component of equation (2-5) is strongly influenced by the shape of the measured objects. SuiteSparse [20] and GRID [21] were developed to improve the computation speed of solving systems of linear equations from FEM. In this paper, a fast frequency-sweeping FEM method with LU decomposition and initial guess/ preconditioning introduced by Lu [22] is used.

### 2.2 Calculation of equivalent complex conductivity

A new method of calculating equivalent complex conductivity of cell suspension along eddy current direction is introduced.

The eddy current density flows in the suspension should be,

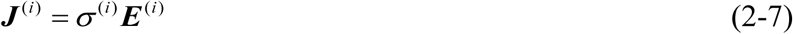

Where ***E***(i) denotes the vector sum of the electrical field on all the edges of each tetrahedral element. *σ*(i) the complex conductivity parameter (with real part the conductivity and imaginary part related to the permittivity) of the of each tetrahedral element.

The model is shown in Figure 3. Assuming there is another background suspension with uniform dielectric and the suspension has exactly the same shape with simulated cell suspension. This uniform suspension is named equivalent model.

Since the normal component of E-field relative to each cylindrical cross-section surface is identical throughout the whole target, then

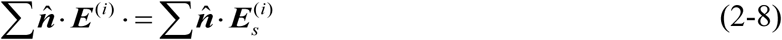

Where, **E**s(i) denotes the background E-field of the of each tetrahedral element (equivalent model); 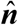 the normal unit vector relative to the surface of target;

Since the equivalent suspension has uniform properties, the electrical background field **E**s is vertical to the cylindrical cross-section surface. Then the equivalent complex conductivity of the original suspension (arranged cells within the suspension) can be deduced from (2-7) and (2-8) that,

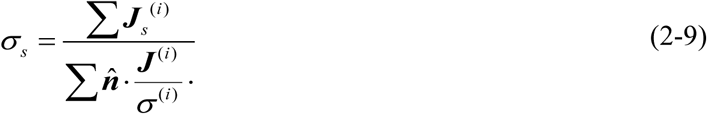

*σ*(i) is the complex conductivity of each tetrahedral element. ***J*_*s*_(i)** is the eddy current density through each cylindrical cross-section of equivalent suspension model. ***J*(i)** is the eddy current density through each cylindrical cross-section of the original suspension model with arranged cells. The current flow direction 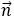 on each cylindrical cross-section surface is shown in Figure 4. The direction can be easily derived:

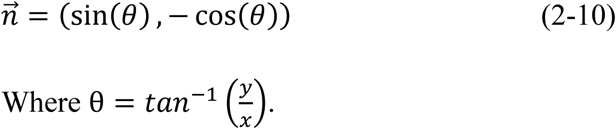

Where 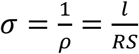.

**Figure 4:**
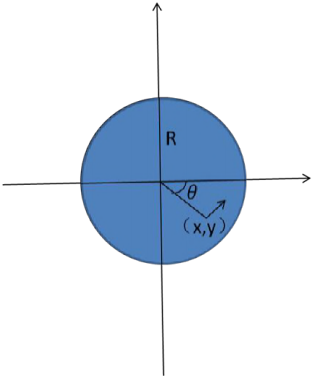
current flow direction (top view of suspension)

Then the complex conductivity could be calculated based on equation (2-9) and the permittivity is simply dividing the imaginary part by jω*ε*_0_ [5].

## 3. Result and discussion

### 3.1 AC conduction contact method - electrode method

#### 3.1.1 Finite element method result

The spherical cell model is shown in Figure 5. The parameters of the cell model are *k*_*m*_ *=* 10^−7^ S/m, *k*_*a*_ *= k*_*c*_ *=* 1 S/m. *k* stands for conductivity. *ε*_*c*_ *= ε*_*a*_ *=* 80, *ε*_*m*_ *=* 5, R=5000 nm, dm=5nm. According to [4] [23].

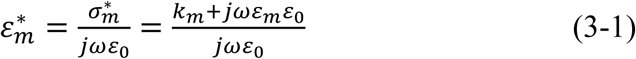

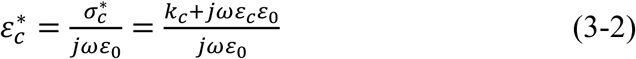

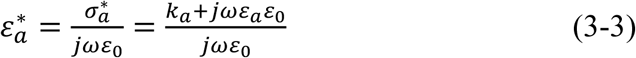

Where *ω =* 2*πf* and *ε*_0_ is the permittivity of vacuum. *f* is the frequency of applied electromagnetic field. *k*_*m*_ is the conductivity of cell membrane. *k*_*a*_ and *k*_*c*_ are the conductivity of extra and intra cellular fluid respectively. *ε*_*m*_ is the permittivity of membrane. *ε*_*c*_ and *ε*_*a*_ are the permittivity of extra and intra cellular fluid respectively. R is the radius of cell, and *d*_*m*_ is the membrane thickness.

**Figure 5:**
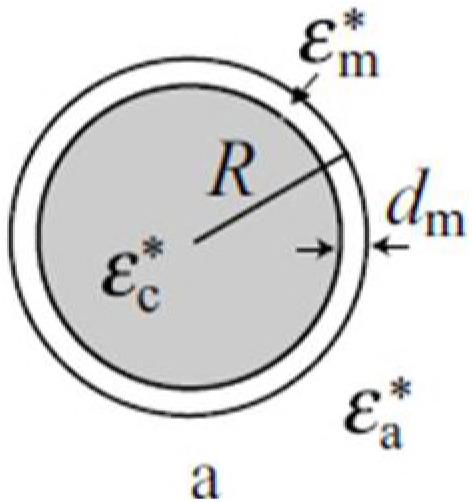
Model of cell

The influence of the shape of the cell is also analysed as followings:

**Figure 6:**
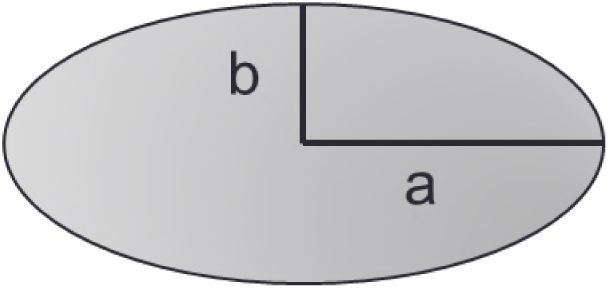
oval cell model

In two-dimension simulation, volume is represented by the area of the cell. The area of a circle is calculated by *S =* π*r*^2^. And the area of an oval is calculated by *S =* πab. Where *a* is the length of the semi-major axis and *b* is the length of the semi-minor axis. So once the radius of cell is confirmed as r, the deformation just need to satisfy one condition *r*^2^ *= ab*.To eliminate the influence of position, all ovals and circles are centred at the same point. And all FEM simulations are carried out under the same meshing accuracy.

Because beta dispersion always occurs during the simulation and only the variation trend of the relative permittivity at low frequency is quite related to the shape of the cell model, a simulation about the cell model deformation is presented. The cell shape parameters are designed following table 1. The cell deform gradually from oval to circle following the parameter of table 1. Since no matter what shape the cell is, it would finally become electrically invisible and the relative permittivity converges to the same value at high frequency as shown in figure 9. So only relative permittivity at low frequency is discussed.

**Table 1:** 2-D deformation parameters

|  |  |  |  |  |  |  |  |  |  |
| --- | --- | --- | --- | --- | --- | --- | --- | --- | --- |
| Deform order | 1 | 2 | 3 | 4 | 5 | 6 | 7 | 8 | 9 |
| Length of a | 2 | 2.4 | 2.5 | 3 | 3.2 | 3.6 | 3.75 | 4 | 4.9 |
| Length of b | 12 | 10 | 9.6 | 8 | 7.5 | 6.67 | 6.4 | 6 | 4.9 |
| Deform order | 10 | 11 | 12 | 13 | 14 | 15 | 16 | 17 | 18 |
| Length of a | 4.9 | 6 | 6.4 | 6.67 | 7.5 | 8 | 9.6 | 10 | 12 |
| Length of b | 4.9 | 4 | 3.75 | 3.6 | 3.2 | 3 | 2.5 | 2.4 | 2 |

**Table 2:** Deformation parameters

| | a $\mu m$ | b $\mu m$ | c $\mu m$ | surface area $\mu m^3$ |
| --- | --- | --- | --- | --- |
| Sphere | 5 | 5 | 5 | 314.0 |
| Scale 1.5 | $5/\sqrt{1.5}$ | $5/\sqrt{1.5}$ | $5*1.5$ | 332.9 |
| Scale 2 | $5/\sqrt{2}$ | $5/\sqrt{2}$ | $5*2$ | 365.8 |
| Scale 3 | $5/\sqrt{3}$ | $5/\sqrt{3}$ | $5*3$ | 434.2 |

**Figure 7:**
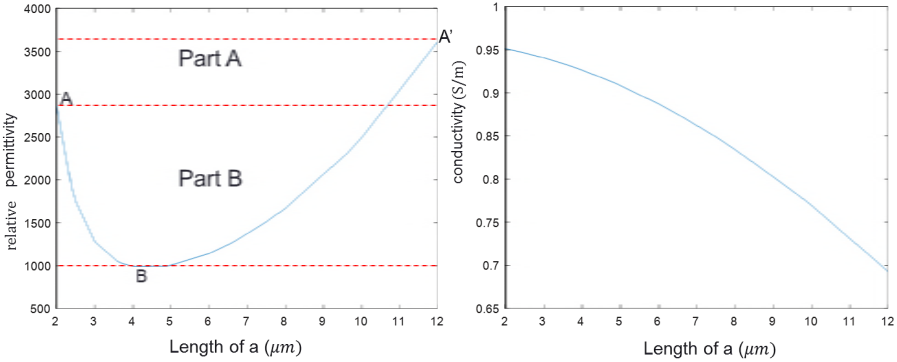
permittivity and conductivity at low frequency(1 kHz)

**Figure 8:**
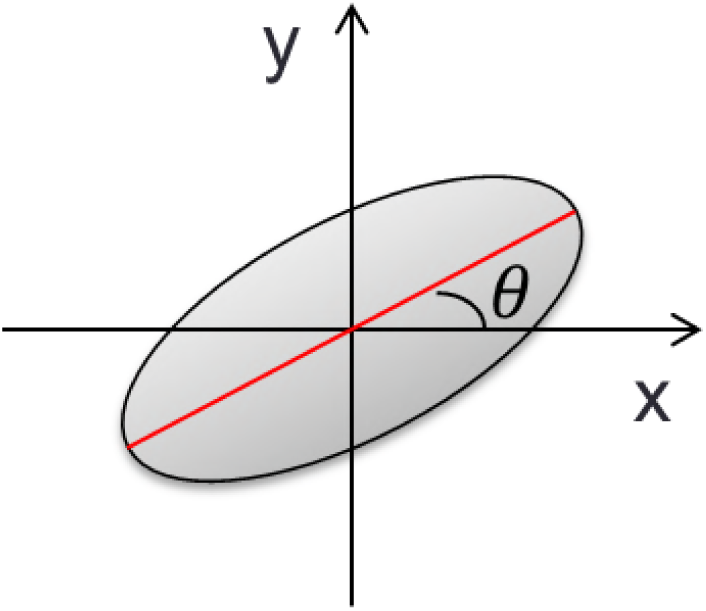
cell orientation model

**Figure 9:**
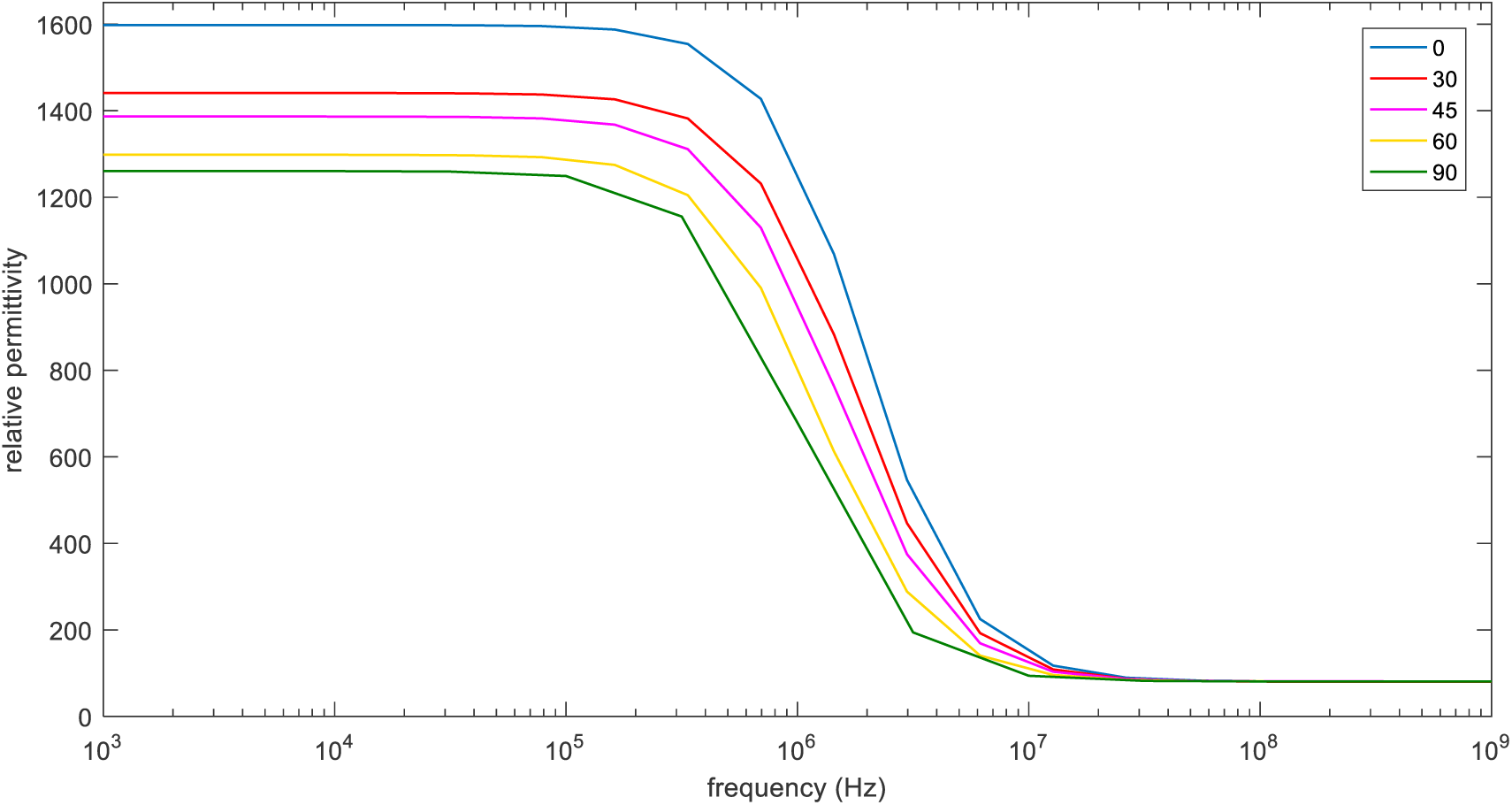
*β-*dispersion with different orientation

As shown in Figure 7, the conductivity decreases monotonically with increasing the length of *a* at low frequency. This is easy to understand as the longer a-axis is the more current will be blocked by the cells. However, the permittivity is not monotonic and not symmetric following the deformation. In part A of figure 7, comparing point A (a=2, b=12) and point A’ (a=12, b=2), the cell models have exactly the same shape parameters but different orientation. So further simulations were carried out to check whether the orientation of cells would influence the permittivity at low frequency.

The cell model rotates from horizontal to vertical and the simulation result is shown in **Error! Reference source not found.** 8. It is obvious that the orientation of the cell would influence permittivity at low frequency. And the smaller *θ* is the larger permittivity will be. This explained the difference between A and A’ in figure 7.

In part B of figure 7, the magnitude of *β*-dispersion has large difference at point A (a=12, b=2) and point B (a=4.9, b=4.9). Asami introduced this phenomenon in his paper [4]. He considered the main factor that leads to magnitude difference of *β*-dispersion was the axial ratio (a/b).

Furthermore, we consider the influence of the membrane thickness (d). To do so, the shape of the cell was kept unchanged and the thickness was increased from 5nm to 10nm.

The internal and external cell fluids are more conductive than the membrane, so the membrane essentially forms a capacitance. Considering the basic function of capacitance:

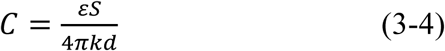

Here the perimeter of the cell can be seen as the surface area S in two-dimension and the membrane thickness can be deemed as the capacitance plate distance d. As shown in figure 10 and figure 11, the permittivity at low frequency decreases while the membrane thickness increases.

**Figure 10:**
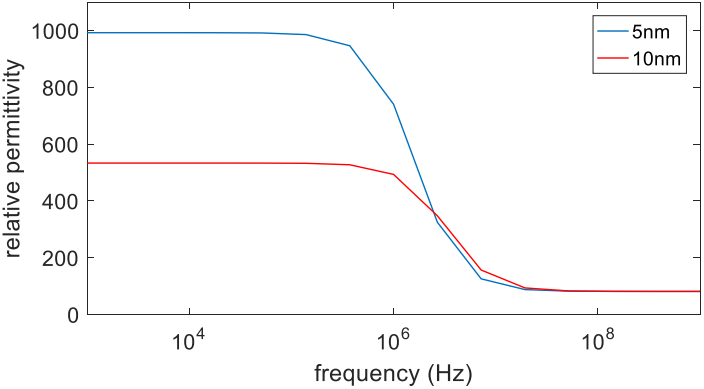
*β*-dispersion of shape Shape: a=4 *µm*, b=6 *µm*

**Figure 11:**
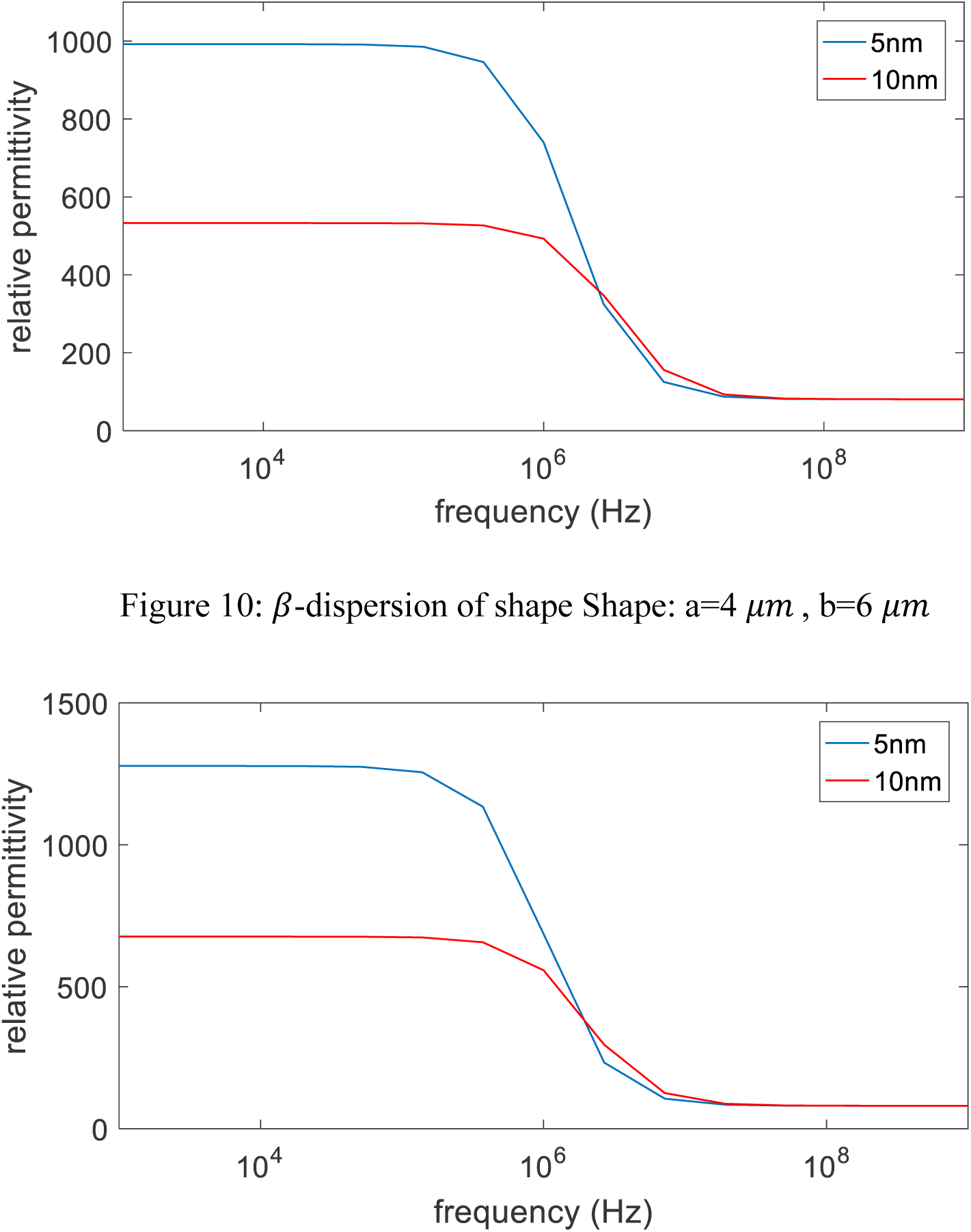
*β*-dispersion of shape Shape: a=3 *µm*, b=8 *µm*

#### 3.1.2 Measurement result

Four-electrode measurement is presented in figure 12. Impedance analyser Solatron 1260 is used to measure the impedance. The effective area of the measurement electrodes is *S =* 2 *cm \** 3 *cm =* 6 *cm*^2^, and the distance between the electric field measuring electrodes is *l = 6.6 cm*. The conductivity could then be calculated by 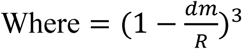. R is the real part of the measured impedance, *ρ* is the resistivity. The measurement samples are saline solution, normal potatoes, and unfreezed potatoes that have been stored in freezer for one day. This measurement is to check the influence of integrity of cell membranes on impedance spectroscopy.

**Figure 12:**
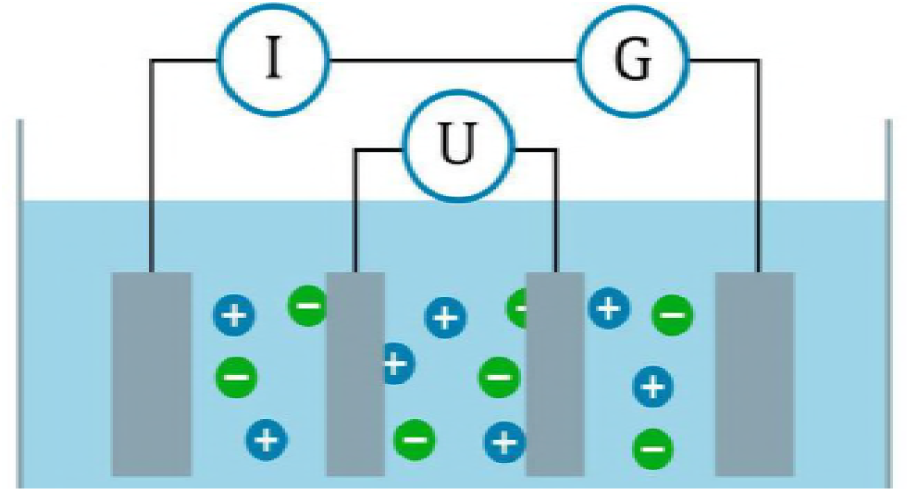
four-electrodes method “https://goo.gl/images/Ama5oF”

The result is shown in figure 13. The measurement result of saline solution shows *α*-dispersion due to the dielectric relaxation at low frequency. Normal potato presents *β*-dispersion starting at around 10 kHz while unfreezed potato shows the similar result with saline solution. This is because the membranes of the potato cells are broken after freezing, the cell membrane no longer blocks current flow and only *α*-dispersion could be observed.

**Figure 13:**
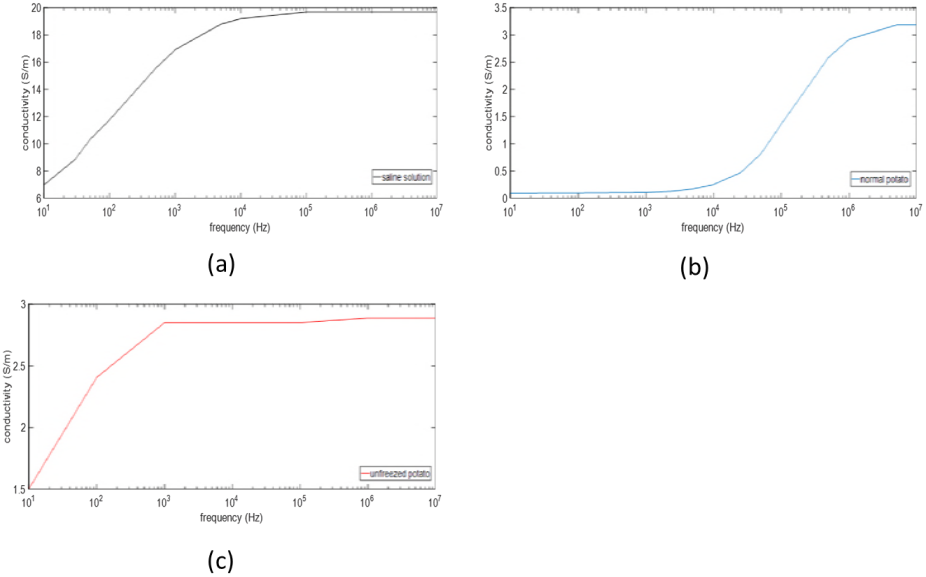
measurement result (a) saline solution with conductivity 4.9 S/m (b) normal potato (c) unfreezed potato

To check the difference between the membranes of normal potato and unfreezed potato, propidium idiode(PI) is used to stain the cells. Since PI stain could not pass the membrane of living cells, it is a reliable method to divide living and dead cells by checking whether the cell is stained. In figure 14, it is obvious that the starch grain inside normal potato cell is not stained while the starch grain is well stained in unfreezed potato cell. This means that the unfreezed cell membrane is broken which is identical with the measurement result.

**Figure 14:**
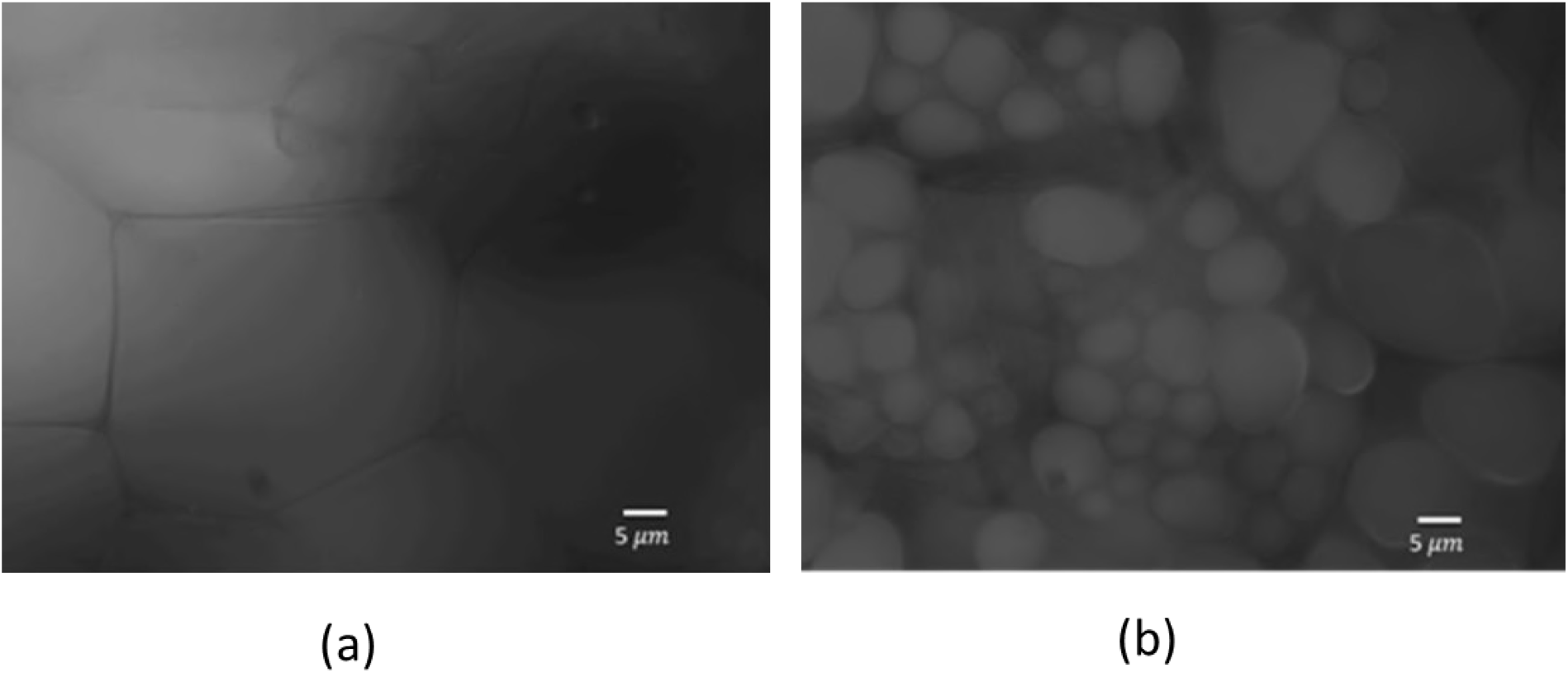
(a) normal potato cell (b) unfreezed patato cell

A simulation about the integrity of cell membranes has been carried out to verify the measurement result. The dead cell model is built by adding break points on the cell membranes as shown in figure 15. Cell models are placed randomly in the simulation in order to simulate real suspension. And the electrical parameters of intra and extra cellular fluid is set to be the same since the cell membranes become permeable after cell death.

**Figure 15:**
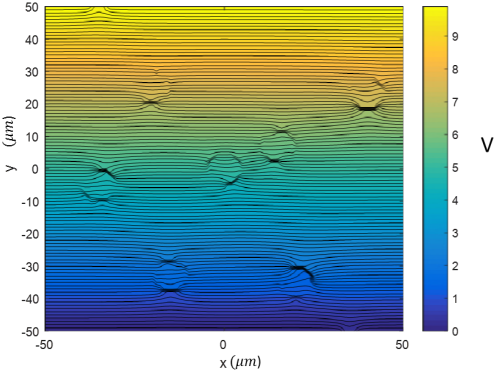
electric potential distribution of break cell model

The simulation result is shown in figure 16. Because *α*-dispersion is caused by ion flows which could not be simulated using finite element method, the simulated result in figure 16 (b) only exhibit a small dispersion. However, the general trend of measurement and simulation result is the same: dead cell suspension has higher conductivity at lower frequency; dispersion of dead cell suspension ends at lower frequency; the conductivity of dead and live cell suspension converges to a steady value at high frequency. This could verify the measurement result about the influence of integrity on bio-impedance.

**Figure 16:**
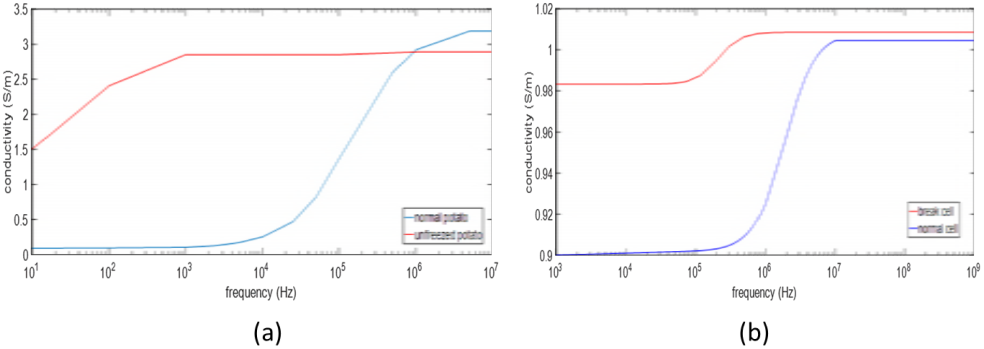
(a) measurement result (b) simulated result

### 3.2 Magnetic induction method

#### 3.2.1 Finite element method result

Figure 17 shows the eddy current distribution under different frequencies. This 3-D simulation shows the same result with 2-D simulation that the cell membrane is not conductive at low frequency [3] and the cell becomes electrically invisible at high frequency since the membrane behaves conductive at higher frequency [15]. Cell models are randomly placed. And the electrical parameters are set to be the same with 2-D simulation.

**Figure 17:**
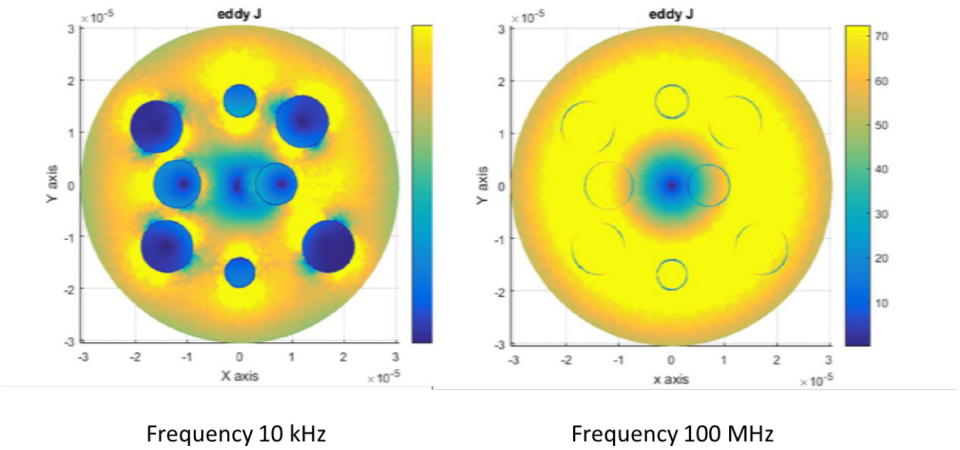
eddy current distribution

The equivalent complex relative permittivity of the cell model in figure 5 can be calculated according to Asami [1]:

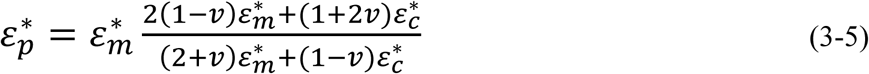

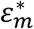, 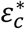 is the complex relative permittivity of membrane and 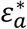 is the complex relative permittivity of cytoplasm. Then the complex relative permittivity of the suspension can be calculated by Wagner’s equation:

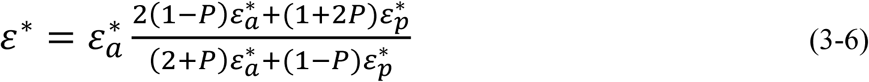

Where P is the volume fraction. 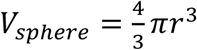 is the complex relative permittivity of the external medium.

**Figure 18:**
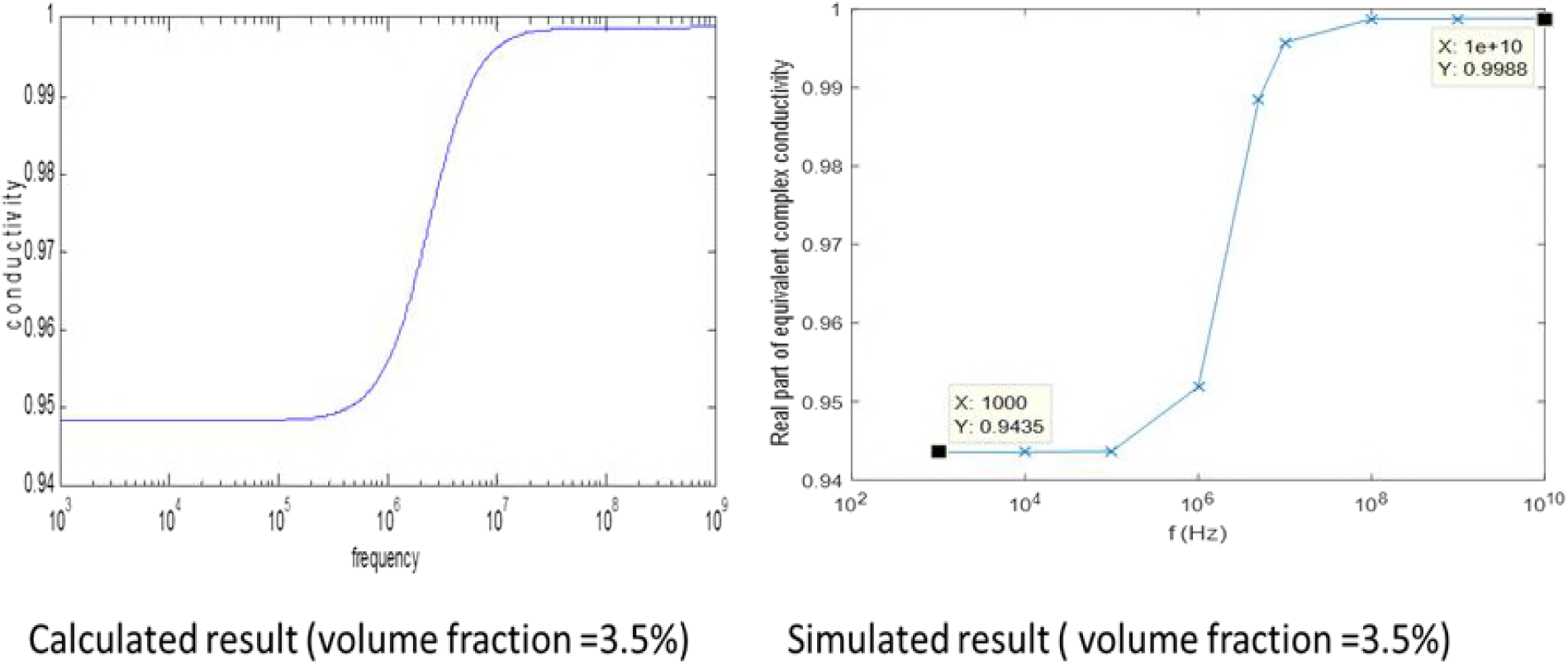
conductivity of analytical solution and FEM simulation

Comparing the FEM simulation result with the analytical solution [4], it can be seen that the magnitude and frequency range of beta dispersion are almost the same. This proves that the three-dimension FEM simulation on cell suspension is feasible and the improved calculation method is correct. In previous 2D simulation, cell shape deformed from circle to oval. And in 3D simulation, cell shape should deform from sphere to ellipsoid. The volume still keeps constant.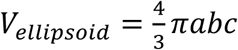 Where a, b and c are the axial length at x, y and z direction respectively. Assume a = b, to keep volume fraction a constant, 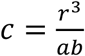

**Figure 19:**
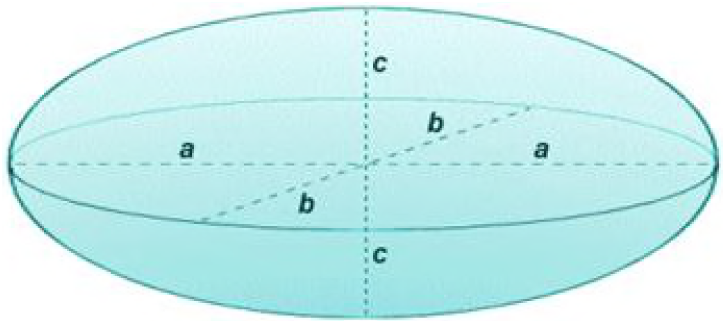
3-D deformation model

**Figure 20:**
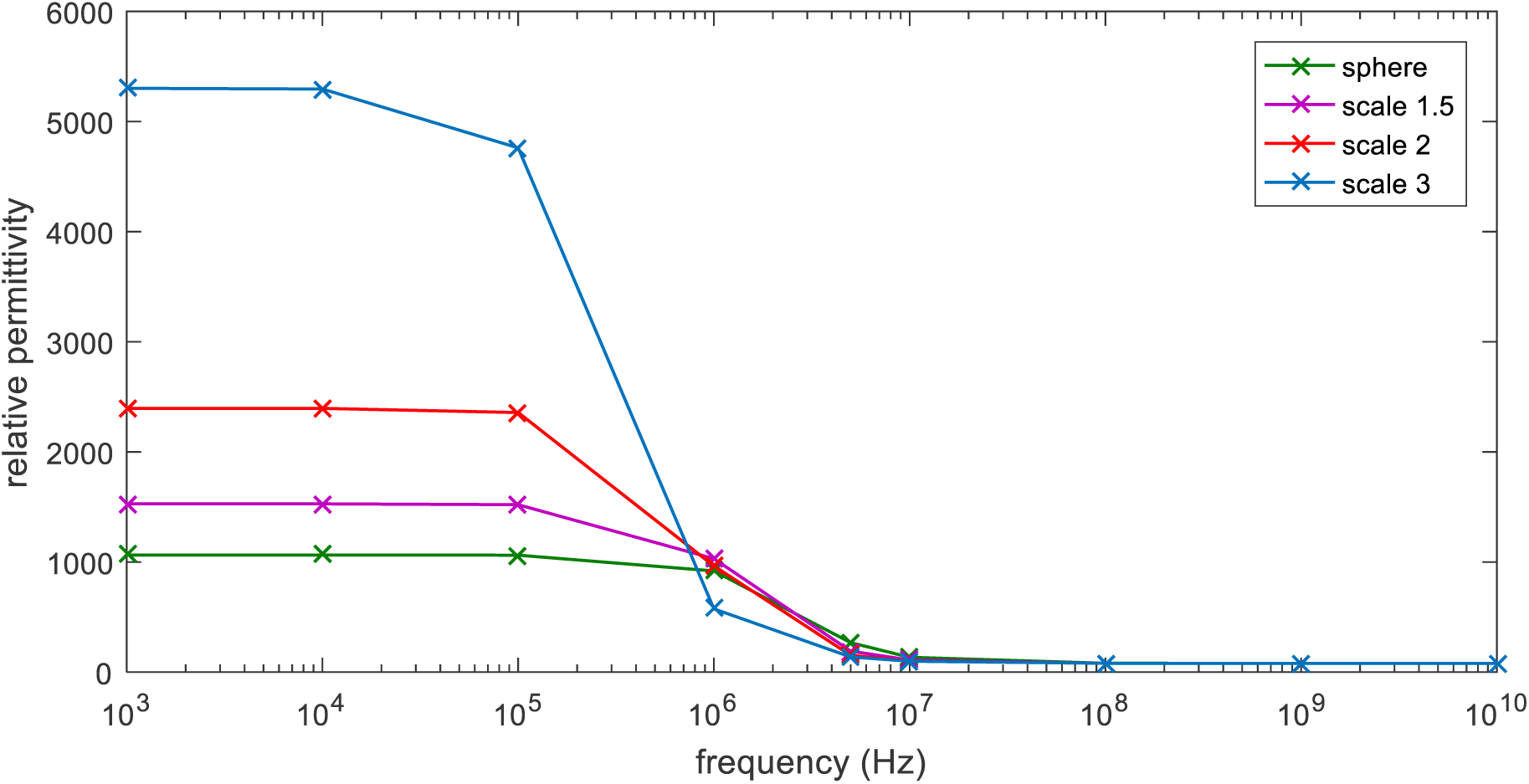
relative permittivity of cell suspension with different deformation shape

The membrane thickness is constant d=5 nm. Surface area (S) increases from sphere to scale 3 ellipsoid. It could be obtained from figure 21 that permittivity reduces with increasing membrane thickness. Because the number of mesh elements is too large (about 1.2 *\** 10^6^), there are only 10 data points of each dispersion.

**Figure 21:**
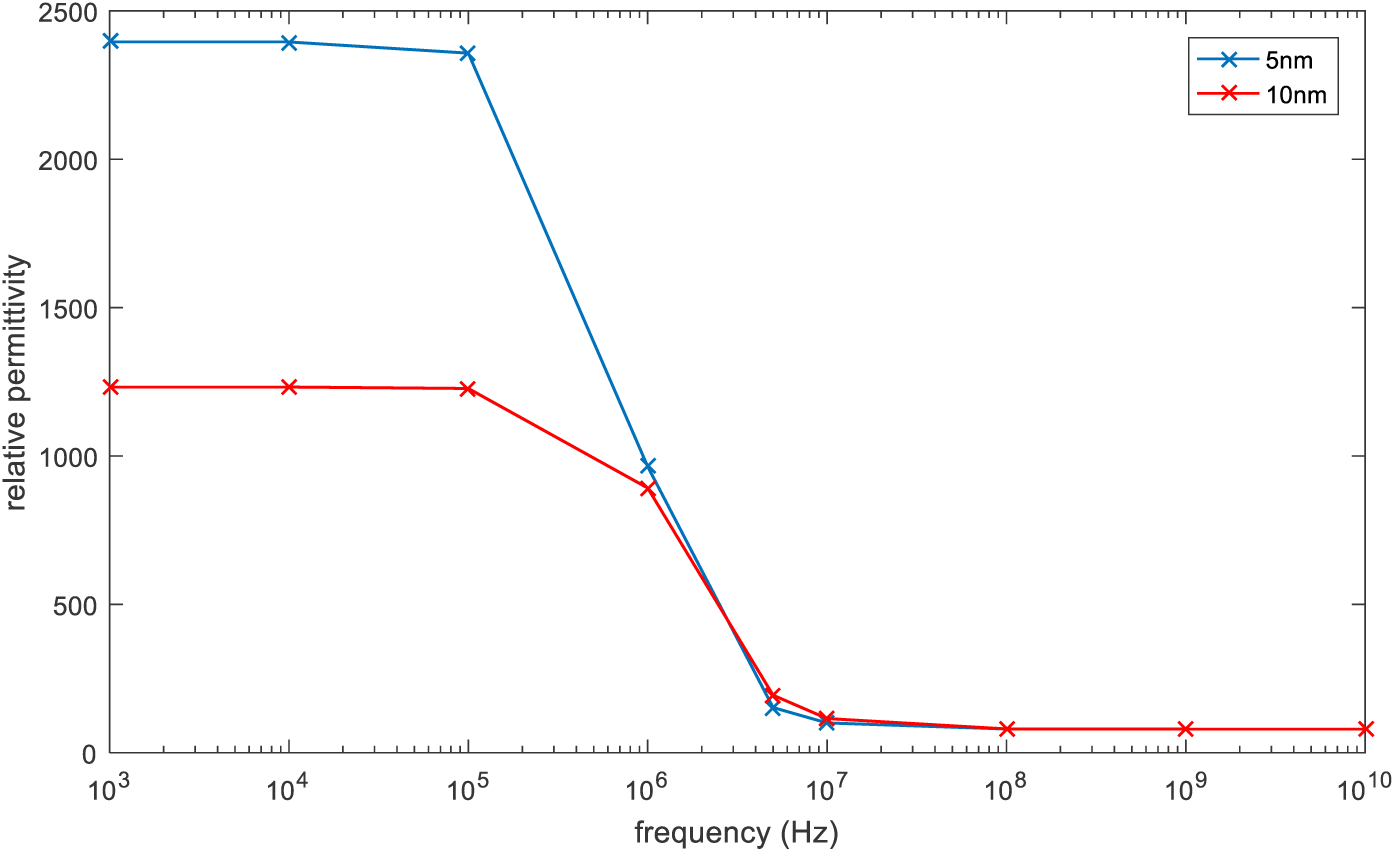
conductivity of parameter scale 2 with different membrane thickness

The 3-D membranes thickness simulation results match the 2-D results which indicate that permittivity reduces with increasing membrane thickness.

**Figure 22:**
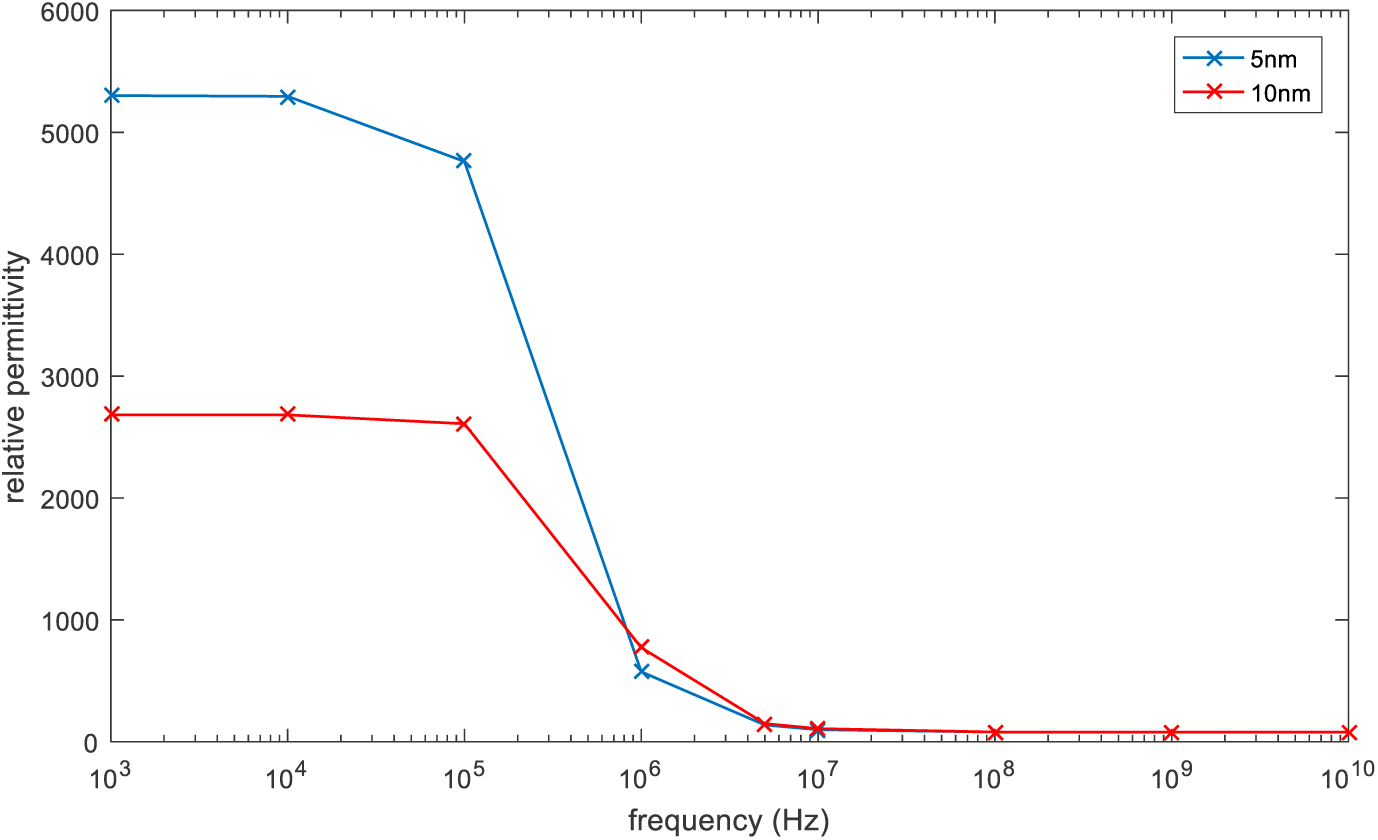
conductivity of parameter scale 3 with different membrane thickness

## 4. Conclusion

Simulations on the influence of cellular structure on bio-impedance are presented in this paper for both electrode method and magnetic induction method. The influence of cell shape and membrane thickness on bio-impedance is investigated and presented. A novel analysis about the membrane thickness influence on bio-impedance spectroscopy and a new method to calculate equivalent conductivity along eddy current direction is introduced. Although the cell models are regularly shaped in this paper, the matlab FEM server could simulate all kinds of irregular cells with different shape and membrane thickness.

The electrode measurement system functioned well on measuring the impedance of saline solution, normal potato, unfreezed potato over a frequency ranging from 10 Hz to 1 MHz. Due to the limitation of hardware, the result is not stable after 1 MHz.

## Acknowledgments

I would like to appreciate and thank my supervisor Dr.Wuliang Yin for his patience, kindness and advices. He helped me a lot not only in academics but also guided me in my life during the one year research.

I would extend my appreciation to my co-supervisor Anthony Peyton for his advices.

Sincerely thanks to Dr Michael O’Toole who helped me a lot and gave me critical advices. And Mr.Mingyang Lu for his kind technical support.

At last, I would thank my parents for their support and encouragement. And thank my girlfriend Yuyang Zhou for her accompany and love.

## Author contributions

J. T. designed experiments and simulations.

J. T. carried out experiments.

J. T. and M. L. carried out simulations.

J. T. analysed experimental and simulation results.

J. T. and W. Y. drafted and revised the manuscript.

J. T., W. Y. and M. L. approved the final version.

